# IL-1R1 dependent signals improve clearance of cytosolic virulent mycobacteria *in vivo*

**DOI:** 10.1101/2020.09.27.315739

**Authors:** Sanne van der Niet, Maaike van Zon, Karin de Punder, Anita Grootemaat, Sofie Rutten, Simone J.C.F.M. Moorlag, Diane Houben, Astrid van der Sar, Wilbert Bitter, Roland Brosch, Rogelio Hernandez Pando, Maria T. Pena, Peter J. Peters, Eric A. Reits, Katrin D. Mayer-Barber, Nicole N. van der Wel

## Abstract

*Mycobacterium tuberculosis* infections claim more than a million lives each year and better treatments or vaccines are required. A crucial pathogenicity factor is translocation from the phago-lysosomes to the cytosol upon phagocytosis by macrophages. The translocation from the phago-lysosome into the cytosol is an ESX-1 dependent process as previously shown *in vitro*. Here we show that *in vivo*, mycobacteria also translocate to the cytosol but mainly when host immunity is compromised. We observed only low numbers of cytosolic bacilli in mice, armadillo, zebrafish and patient material infected with *M. tuberculosis, M. marinum* or *M. leprae*. In contrast, when innate or adaptive immunity was compromised, as in SCID or IL-1R1 deficient mice, a significant number of cytosolic *M. tuberculosis* bacilli were detected in lungs of infected mice. Taken together, *M. tuberculosis* infection is controlled by adaptive immune responses as well as IL-1R1-mediated signals that result in clearance of cells containing cytosolic mycobacteria *in vivo*.

**Importance:** For decades, *Mycobacterium tuberculosis* is one of the deathliest pathogens known. Despite infecting approximately one third of the human population, no effective treatment or vaccine is available. A crucial pathogenicity factor is the subcellular localization, as *M. tuberculosis* can translocate from the phago-lysosome to the cytosol in macrophages. The situation *in vivo* is more complicated. In this study we establish that high level cytosolic escape of mycobacteria can indeed occur *in vivo*, but mainly when host resistance is compromised. The IL-1 pathway is crucial for the control of the number of cytosolic mycobacteria. The establishment that immune signals result in clearance of cells containing cytosolic mycobacteria, connects two important fields: cell-biology and immunology which is vital for the understanding of the pathology of *M. tuberculosis*.

## Introduction

*Mycobacterium tuberculosis* (*Mtb*) is not only one of the most deadliest pathogens in history, but it continues to claim an estimated 1.5 million human lives per year (1) and is a big threat for the future as multidrug resistant strains are arising with no effective treatment nor vaccine available. The treatment success rate is just 56% and in approximately 6.2% of the cases, infection is caused by extensive drug resistant *Mtb* (1). Furthermore, *Mtb* is the leading cause of death among HIV infected patients. In patients with AIDS, tuberculosis can thrive because their immune system is impaired by CD4^+^T cell loss, which secondarily affects many other immune compartments (reviewed in (2)).

For an effective response to *Mtb* infections, both the innate and the adaptive immune system are important. The development of an active pulmonary *Mtb* infection is related to a disordered immune balance, which results in the inability of the host to keep the infection under control (3). The first immune response is the innate response (reviewed in (4)) including a series of cells that come in contact with *Mtb* such as alveolar macrophages. Alveolar macrophages provide a nutritionally permissive niche (5) and are critical for dissemination of the bacteria in the lung, spreading the infection from the alveoli to the interstitium (6). Here *Mtb* infects other cell types like neutrophils, monocyte derived macrophages and dendritic cells (DC). Since DCs present antigens via MHC class I and II to T cells, they function as a connection between the innate and adaptive immune system (7, 8). Upon antigen presentation to T cells, CD4^+^T cells produce IFNγ, which is involved in the enhancement of macrophage killing and plays an important role in granuloma formation (9). Indeed in mice (10) and humans (11) loss of IFNγ or its receptors, acting as single non-redundant factor, leads to TB disease. In order to spread to new individuals, *Mtb* needs to cause pulmonary lesions (12). Inflammation driven by IL-1 contributes to host resistance to *Mtb* (13, 14). Mice that lack the IL-1R1, IL-1α or IL-1β display high susceptibility to *Mtb* infection (15) with uncontrolled bacterial replication in the lungs, again demonstrating a non-redundant role of one key host pathway.

We hypothesized that bacterial translocation from the phagosome to the cytosol might also be regulated by IL-1. *In vitro, Mtb* can translocate from the phago-lysosome to the cytosol (16–23) in an ESX-1 dependent manner (16, 21, 22). This system is responsible for the secretion of a number of proteins, including EsxA (ESAT-6) and EsxB (CFP-10) (reviewed in (24)). When this secretion system is not present, as is the case for *M. bovis* BCG, cytosolic localization is abrogated in *in vitro* macrophage systems, rendering the bacteria restricted in a membrane enclosed phago-lysosome. Reintroducing the extended *esx-1* locus in BCG allowed the translocation to the cytosol and increased virulence providing clear evidence for an essential role of this Type VII secretion system in escape (16, 25–28). In addition, it is shown that virulent *Mtb* can form cords in the cytosol and not in the phagosome in human lymphatic endothelial cells *in vitro* (23). This cording is dependent on the ESX-1 secretion system and PDIM glycolipids. The formation of cords in the cytosol rather than in phagosomes suggests a permissive environment for bacterial replication in the cytosol. When *Mtb* is present in the cytosol its bacterial DNA is sensed by cyclic guanosine monophosphate-AMP synthase (cGAS) (29–32). This detection is dependent on the presence of a functioning ESX-1 system, suggesting that it is dependent on pathogen-induced cytosolic localization. Cytosolic bacteria co-localize with cytosolic ubiquitin, while this is not the case when the mycobacteria are present in the phagosome (16). Another factor involved in the cytosolic localization is Rv3167c, which regulates the escape of *Mtb* from the phagosome, since a mutant unable to produce this protein (MtbΔRv3167c) displayed increased cytosolic escape (33). Other virulent factors involved in escape from the phagosome are PDIM glycolipids located on the outer membrane of *Mtb* (19, 25, 34). Mycobacterial strains that lack PDIM are less capable of damaging the phagosomal membrane, resulting in less *Mtb* in the cytosol of THP-1 macrophages. Recently it was shown that phagosomal rupture causes activation of NLRP3-dependent IL-1β release and pyroptosis, a programmed form of cell death (35) facilitating the spread of bacteria to other cells.

While most studies focused on *in vitro* experiments using cultured macrophages, the sub-cellular localization of *Mtb* and the factors affecting cytosolic localization and pathogenesis are less intensively studied *in vivo*. When macrophages purified from bronchoalveolar lavages from TB infected patients were analyzed by electron microscopy, it was found that *Mtb* is primarily localized in phagosome-like compartments (36, 37). The phagosomal localization does not affect the ability of *Mtb* to proliferate, and it is long known that when *Mtb* is located in phagosomes it is able to arrest its maturation (38, 39) up to 5-7 days (40). More recently, it was shown *in vivo* that *Mtb* is able to translocate to the cytosol as early as three hours post infection using a Förster resonance energy transfer (FRET)-based detection system (41). This study also demonstrated that the pH of the lysosomes plays a role in the cytosolic localization; when the lysosome is more acidic, less *Mtb* is present in the cytosol at 3 days post infection (41).

To examine whether activation via adaptive or innate immunity pathways would affect the capability of mycobacteria to translocate into the cytoplasm *in vivo*, we tested the subcellular localization of different mycobacterial species in zebrafish, armadillo and mice models as well as in patient material. In adult zebrafish and zebrafish embryos we used *M. marinum*, a close homologue of *Mtb* that is also known to escape the phago-lysosome in an ESX-1 dependent manner (42). *M. leprae* is also known to escape to the cytosol (22), probably using a similar mechanism, although some of the members of ESX-1 system (like esxC, esxG, esxS) are pseudogenes, it has functional ESAT-6 (esxA) and CFP-10 (esxB), the two most important components needed for cytosolic escape. We used both skin biopsies of leprosy patients and the armadillo model for *M. leprae* since, the armadillo model is known to exhibit the entire clinical spectrum of leprosy (43). For *Mtb* both SCID mice that lack both T and B cells, and IL-1R1 knockout mice were compared to determine their ability to limit cytosolic escape *in vivo* in infected cells. Here we show that while cytosolic localization is limited by both innate and adaptive immunity, IL-1 seems to be a key effector pathway in controlling cytosolic translocation of *Mtb*.

## Results

### The pH of the phagosome and lysosome does not affect cytosolic localization of *M. marinum*

*Mtb* blocks maturation and acidification of the phago-lysosome, promoting its intracellular survival (38, 39, 44–46). Phagosomal acidification is essential for increased activity of the lysosomal digestive process and thus for degradation of its content (47). *Mtb* partially avoids acid mediated killing by blocking fusion between lysosomes and phagosomes (36, 39) and the secretion of antacid known as 1-tuberculosinyladenosine (TbAd) (48). In addition, it is shown that the mycobacterial cell wall plays a role in the resistance to acidic environments (reviewed in (49)). We hypothesized that cytosolic escape is a fourth mechanism to avoid lysosome mediated killing. To determine whether these mechanisms are interdependent, we examine if the phagosomal pH affects translocation from the phagosome to the cytosol. To exclude the effect of TbAd, we utilized *M. marinum*, which does not express TbAd (50) in THP-1 cells. The acidity of the phagosome and the lysosome was measured using lysotracker and by incubation with (N-(3-((2,4-dinitrophenyl)amino)propyl)-N-(3-aminopropyl) methylamine (DAMP), a weak basic amine that will be taken up by acidic organelles in live cells (36). After fixation and sample preparation, the DAMP was visualized by Transmission Electron Microscopy (TEM) using immuno-gold labelling with αDNP antibody conjugated to a gold particle. The more acidic the phagosome or lysosome, the more DAMP was present, thus resulting in a higher label density in acidic organelles. As expected, upon *M. marinum* infection of THP-1 cells, low amounts of DAMP labelling were observed surrounding cytosolic *M. marinum* (Fig. S1A/1A’) while more labeling was detected in *M. marinum* containing phagosomes (Fig. S1A/1A”). We next blocked acidification with 10nM Concanamycin B (ConB), an inhibitor of vacuolar ATPases which prevents acidification of endosomes and lysosomes. When cells were treated with ConB and infected with *M. marinum*, both a lower label density was measured and the lysotracker imaged with fluorescence microscopy confirm that the pH is less acidic in the phagosome/lysosome when treated with ConB as already well described (36, 51) (Fig. S1B). We next examined whether a raising pH would affect the translocation efficiency of *M. marinum* to the cytosol. The percentage of cytosolic bacteria in THP-1 cells treated with ConB or no treatment control was determined both at 24h and 48h after infection with *M. marinum*, which is the known timeframe for escape (16) (Fig. 1A,B). The number of bacteria present in CD63 labeled compartments (phago-lysosomal), membrane enclosed but not CD63 positive compartments (phagosomes) and the number of bacteria in the cytosol were counted (Fig. S2A). At both 24h and 48h after infection no difference in the percentage cytosolic bacteria was detected for untreated or ConB treated cells. This indicates that a higher pH has no effect on *M. marinum* translocation to the cytosol. In conclusion, a raised lysosomal pH has no effect on the percentage of bacteria translocating to the cytosol in THP-1 cells.

**Legend Fig 1.**
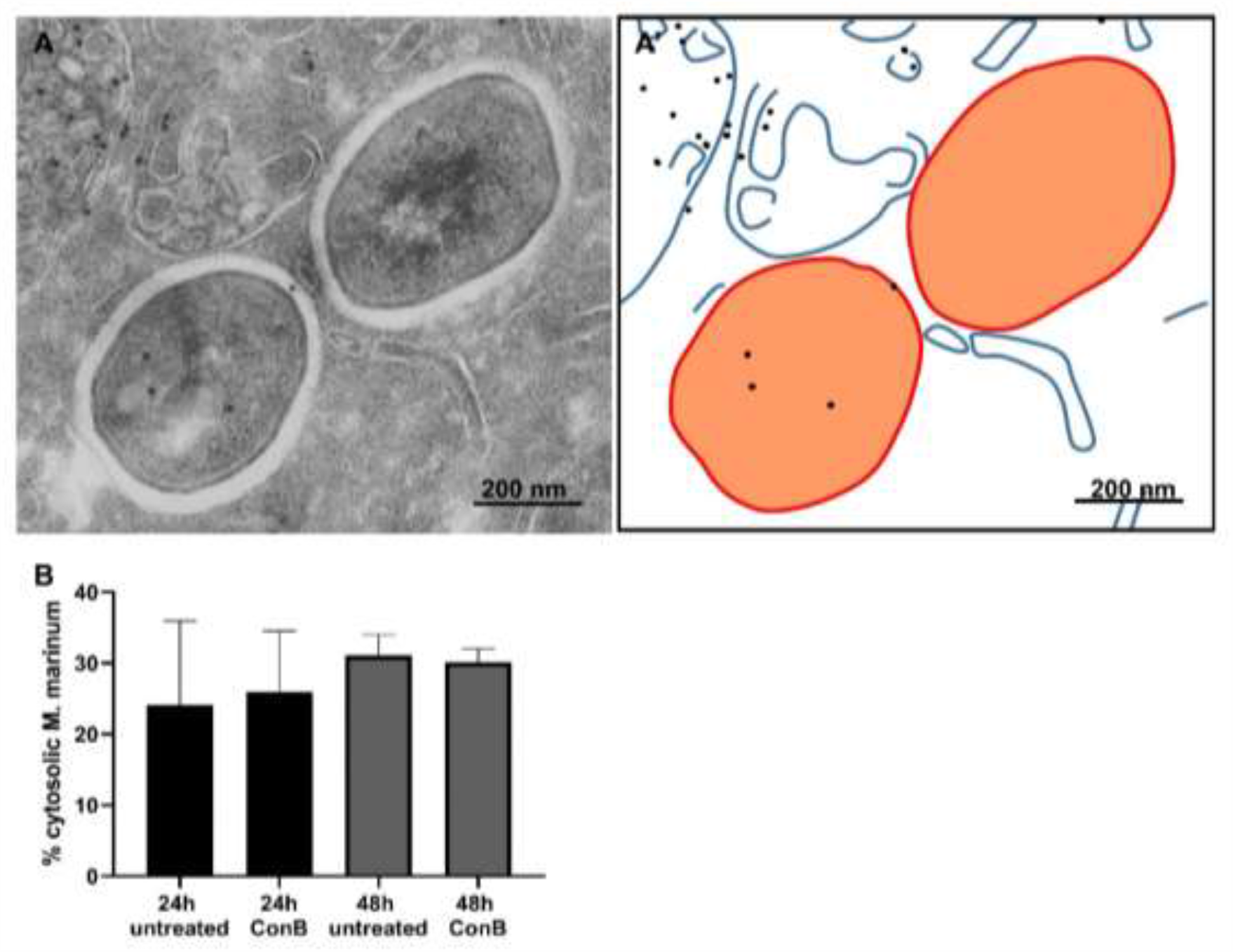
The pH of the phago-lysosome does not affect cytosolic translocation of *M. marinum*. A) Electron micrograph of a THP-1 cell infected with *M. marinum* for 24 hour in the presence of ConB showing *M. marinum* in the cytosol without membrane and CD63 labeling. CD63 immunolabelling indicated by 10 nm gold particles is present on the multivesicular lysosome in the left top corner. A’) schematic representation of micrograph in A with in blue lines host membranes, black dots CD63 labelling indicated by 10 nm gold particles, orange bacteria and bar represents 200 nm. B) Quantification of the percentage of *M. marinum* in the cytosol 24 and 48 hours after infection using immuno-gold labelling for CD63 (see also Fig. S2A). Cells were treated with ConB to raise lysosomal pH, which did not affect the percentage of cytosolic *M. marinum*. Error bars represent the standard deviation of 3 experiments.

### *M. marinum* translocates to the cytosol in embryonic but not adult zebrafish

After studying the effect of the pH on the ability of *M. marinum to* translocate to the cytoplasm *in vitro*, we next focussed on the subcellular localisation *in vivo*. To address whether either adaptive or innate immunity was required to limit bacterial cytosolic escape we used a zebrafish model combined with *M. marinum* infection. Among other age-related differences, zebrafish larvae do not have a fully developed adaptive immune system in contrast to adult zebrafish (52). Zebrafish embryos and adult zebrafish were infected with *M. marinum* or *M. marinum* Tn::ESX5 mutant and tissue was fixed for TEM analysis. For analysis of adult zebrafish a specific *M. marinum* Tn::ESX5 mutant was used as this infection was shown to cause hypervirulence (53) and thus large amounts of bacteria can be detected *in vivo* using TEM. The following conditions were analysed using TEM: whole zebrafish embryo day 9 and the spleen of adult zebra fish at day 11 (Fig. 2A,B). In zebrafish embryos, we have previously demonstrated that injected *M. marinum* with fluorescent tags is present in phagocytic cells in close proximity of blood vessels and endothelial cells of these blood vessels (54). Here we injected the zebrafish with untagged *M. marinum* and detected bacteria in similar phagocytic and endothelial cells, close to or part of the blood vessels and in phagocytic cells spread through the tissue. In these cells, the percentage of cytosolic *M. marinum* was determined by counting the number of mycobacteria in a membrane enclosed compartment and in the cytoplasm. Discrimination between phagosome and phago-lysome based on immuno-gold labelling with lysosomal markers available in humans or mice (CD63, LAMP1) was not possible as no lysosomal nor cellular markers are available for immuno-gold labelling in zebrafish. Alternatively, actin antibodies were used as a cellular cytoskeleton marker. In embryos infected for 9 days 32% of *M. marinum* was present in the cytosol (Fig. 2C and Fig. S2B). Noteworthy, ESX-1 mutants remain restricted in the lumen of the blood vessels and not able to pass the basal lamina of the blood vessel wall (54). As only part of the ESX-1 mutant bacteria is intracellular and the majority restricted in the lumen of the vessels, the ESX-1 mutants were not included in this study. Moreover, there is ample literature on the ESX-1 localisation in membrane enclosed compartments (16, 22, 25–28, 35). Out of the four adult zebrafish analyzed, one was discarded as no bacteria could be deteced. In the other three fish ample bacteria were detected and catagorized as phagosomal or cytosolic based on the presence of membrane. In adult zebrafish less than five percent of the bacteria were found in the cytosol (Fig. 2C). Thus, adult but not embryo zebrafish were able to limit bacterial cytosolic escape. This outcome suggests the presence of an adaptive immune system promotes bacterial containement in the phago-lysosome, although other age related factors could be at play.

**Legend Fig 2:**
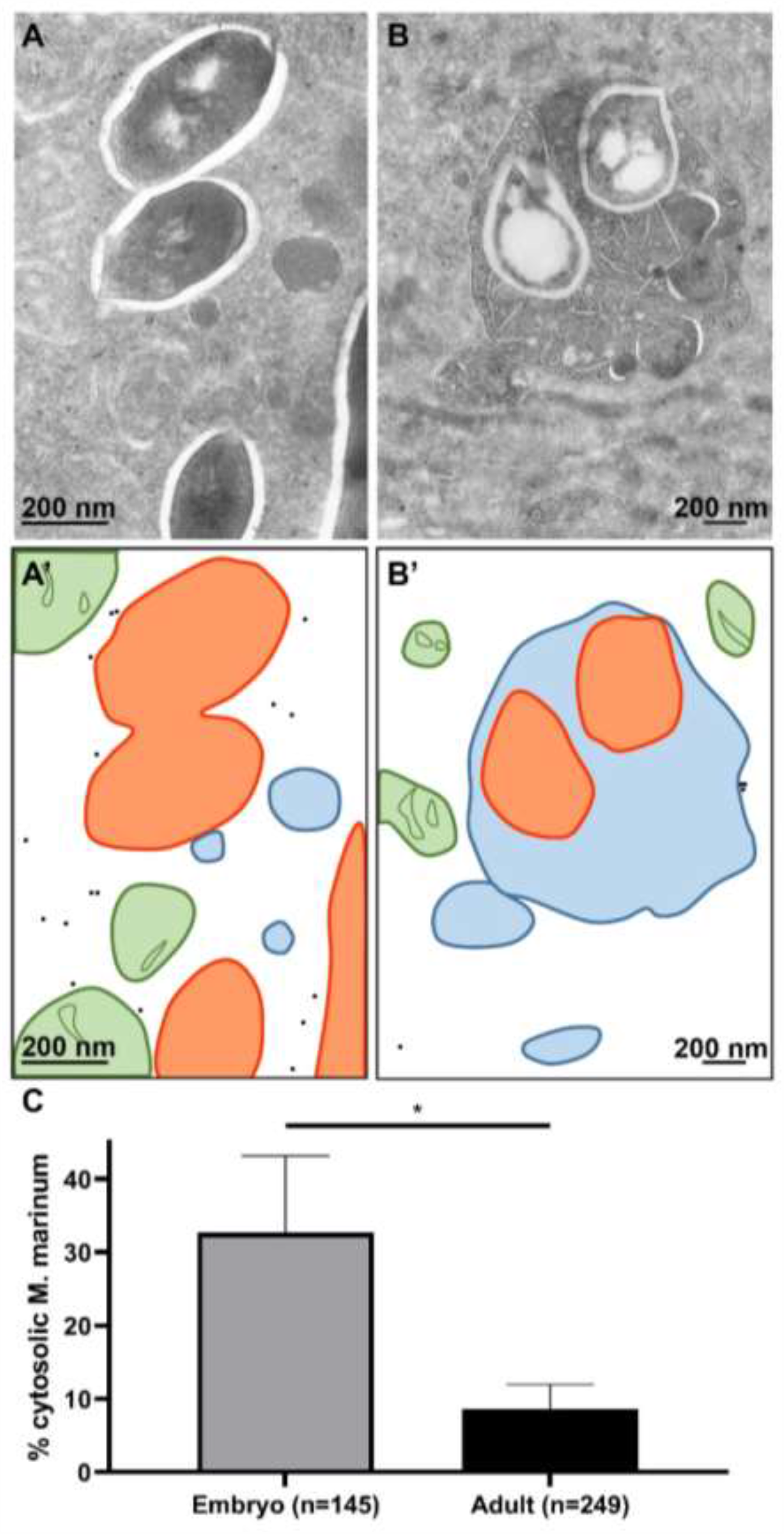
Cytosolic localization of *M. marinum* in zebrafish is abundant when the adaptive immune system is not yet developed. Embryo and adult zebrafish infected with *M. marinum* were analysed using TEM. A) Cytosolic *M. marinum* in zebrafish embryo tissue. B) Phagosomal *M. marinum* ΔESX5 in adult zebrafish tissue. A’) and B’) Schematic representation of A and B, with black dots indicating actin immuno-gold labelling, orange lines *M. marinum* and blue lines phagosomal and host membranes, green lines mitochondria. C) Quantification of the percentage of cytosolic *M. marinum* at embryo day 9 and in spleen adult zebrafish at day 11 (see also Fig. S2B). Error bars indicate standard deviation between 3 different zebrafish embryos and 3 adult fish, n represents the total number of bacteria counted.

### Cytosolic localization of *M. leprae* is restrained in both armadillo and patient skin

To examine whether cytosolic localization can only be detected in early, innate stages of infections, we used another model organism; *M. leprae*, the causative agent for leprosy. Like *Mtb*, these bacteria have been shown to translocate to the cytosol in an *in vitro* model (22). We studied early (day 3) and late stage (day 21) infections in armadillos and in addition, skin biopsies were taken from 4 lepromatous leprosy (LL) patients with a well-established infection. The skin of the abdomen of armadillos was infected with both unviable (irradiated) as well as viable *M. leprae*. At the site of infection, loss of pigment was visible which increased during infection progression (Fig. 3A and B). No differences were observed in loss of pigment between the sites infected with irradiated *M. leprae* and viable *M. leprae*. At the border of pigment loss, armadillo skin biopsies infected with viable *M. leprae* were fixed for TEM analysis, to determine if the localization of the bacteria is cytosolic versus phagosomal. Immuno-gold labelling against *M. leprae* specific anti-Cell Wall Protein was used to verify that these are indeed mycobacteria. Similar to the localization of *M. marinum* in adult zebrafish, the majority of the bacteria were present in membrane enclosed phagosomes and cytosolic bacteria were only occasionally detected at both day 3 (Fig. 3C) and 21 after infection. In addition, skin biopsies of 4 different lepromatous leprosy patients taken from the border of the infection as defined by the depigmentation line and were analyzed using TEM and immuno-gold labelling for *M. leprae* specific anti-Cell Wall Protein and lysosomal markers such as Cathepsin D (Fig. 4), LAMP1, and CD63. Worth mentioning is that based on the ultrastructure of the cells, and more specifically the ultrastructure of the cell nucleus or the presence of multiple lysosomes, most bacteria reside in macrophage-like cells. From the 4 patients, individual bacteria were detected and classified for their subcellular localization based on the presence of surrounding membrane and 2 or more gold particles detecting Cathepsin D (Fig. S2C). Similar to the armadillo and adult zebrafish, the percentage of cytosolic mycobacteria was low (1.1%), while over 700 bacilli were assessed. In conclusion, in both armadillo and human skin, a low percentage of cytosolic *M. leprae* bacteria is present at all measured stages of infection.

**Legend Fig 3:**
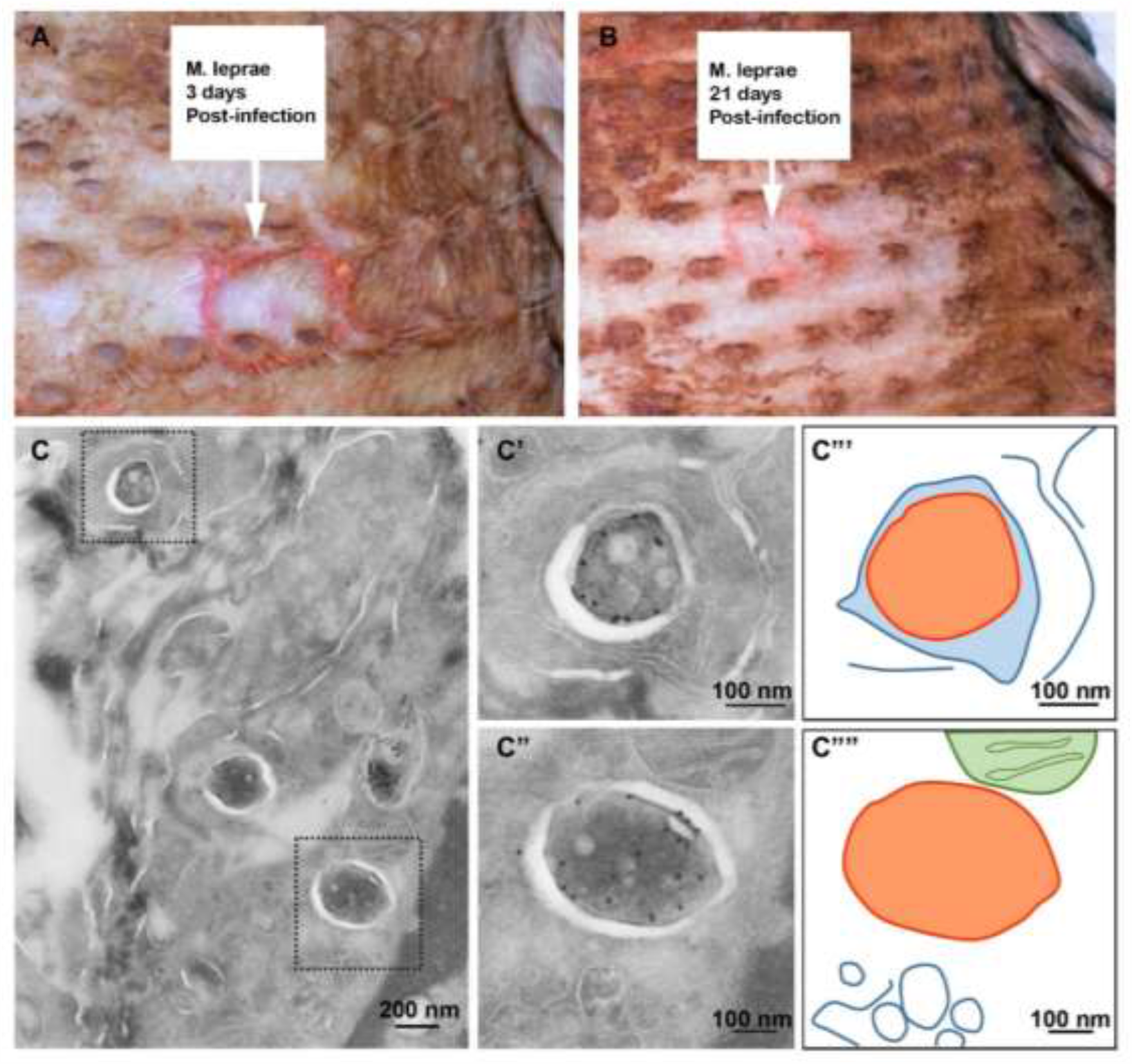
Restrained cytosolic *M. leprae* localization in armadillo skin biopsies. Live *M. leprae* injected in the skin of the armadillo at the red circle indicated by the white arrow text box. The skin was observed at A) 3 days after infection and B) 21 days after infection. At infection sites loss of pigmentation was detected. C) TEM image of infected armadillo skin biopsy 3 days after infection with viable *M. leprae*. Immuno-gold labelling using αCWP to indicate *M. leprae*. C’) enlargement of boxed area in C, *M. leprae* enclosed by host membranes and thus phagosomal. C”), enlargement of lower boxed area in C, *M. leprae* was not enclosed by host membranes and thus cytosolic. C’”), C’”‘) Schematic representation of C’ and C” with in orange *M. leprae* and the blue lines indicate host membranes and green mitochondrial membranes. Total number of bacteria detected at day 3 is 47 and at day 21 is 45.

**Legend Fig 4.**
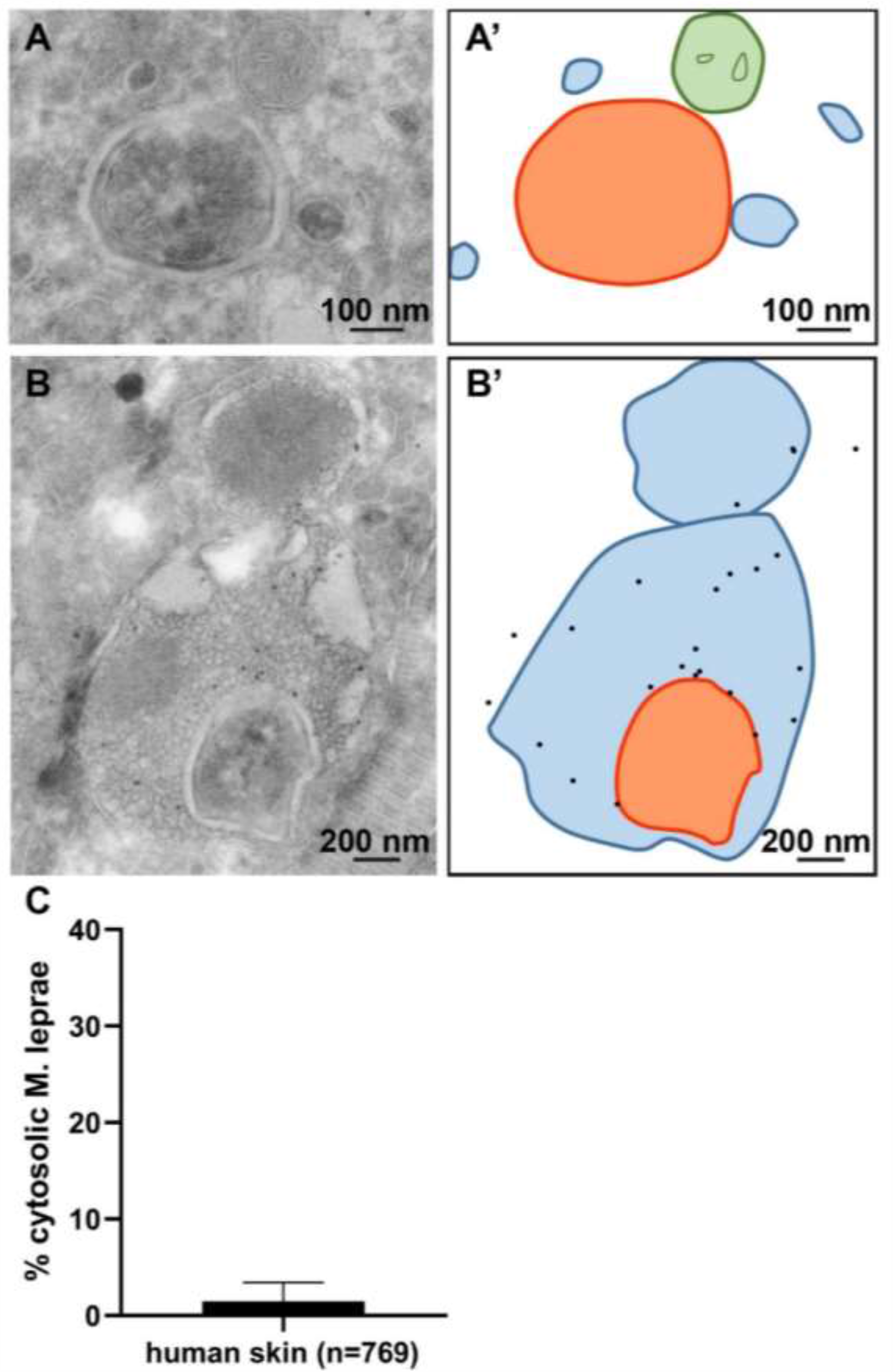
Restrained cytosolic localization of *M. leprae* in human skin biopsies. Leprosy patient skin biopsies were analyzed using TEM. Immuno-gold labelling for Cathepsin-D was used to label lysosomes and phagolysosomes. A) TEM image of patient skin biopsy with cytosolic *M. leprae*, the mycobacteria was not enclosed with host membranes. A’) Schematic representation of A, orange indicate *M. leprae*, blue lines indicate host membranes, and green mitochondria. B) *M. leprae* present in the phagolysosome, the mycobacterium is enclosed by host membranes, immunogold labelled for lysosomal marker Cathepsin-D. B’) Schematic representation of B, black dots indicate Cathepsin-D labelling, orange indicate *M. leprae* and the blue lines indicate phagolysosomal membranes. C) Quantification of the average percentage of *M. leprae* present in the cytosol, error bar indicates standard deviation from 4 different patients, n represents the total number of intracellular bacteria (see also Fig. S2C).

### Immunocompetent BALB/c, but not SCID mice can contain *M. tuberculosis* inside phagosomes of infected pulmonary cells

After showing that neither *M. leprae* nor *M. marinum* translocate in high numbers to the cytosol even at later stages of infection, we next wanted to study the subcellular localization of *M. tuberculosis* longitudinally in mice. To do so BALB/c mice were infected with the virulent *Mtb* strain H37Rv and lung tissue was fixed for EM analysis at day 2, 7, 21, 45 and 120 after infection. Based on the ultra-structure of the nucleus and/or the high number of lysosomes, the intracellular bacteria were detected in macrophage-like cells. These samples were immunogold labelled for lysosomes using LAMP1 or Cathapsin D and imaged using TEM (Fig. 5). As for *M. marinum* in THP1 cells and *M. leprae* in skin, the number of bacteria present in LAMP-1 labeled phago-lysosomes, membrane enclosed but not LAMP-1 labelled phagosomes and the number of bacteria in the cytosol were counted (Table 1S). We found that cytosolic localization was highest at day 7 of infection, with still less than five percent of *Mtb* detectable in the cytosol and most bacteria residing in a membrane enclosed compartment. To determine if patient derived strains behave similar to H37Rv, BALB/c mice were infected with two additional *Mtb* strains from lineage 2, often referred to as Beijing family *Mtb* (1998-1500 ancient Beijing) and multi drug resistant strain (2002-0230 Beijing). EM analysis was done at 21, 45 and 120 dpi and demonstrated also low amounts cytosolic bacteria (Table S1). Thus, as shown previously in immune-competent mice, different strains of *Mtb* reside primarily inside membrane enclosed compartments and not inside the cytosol.

**Legend Fig 5.**
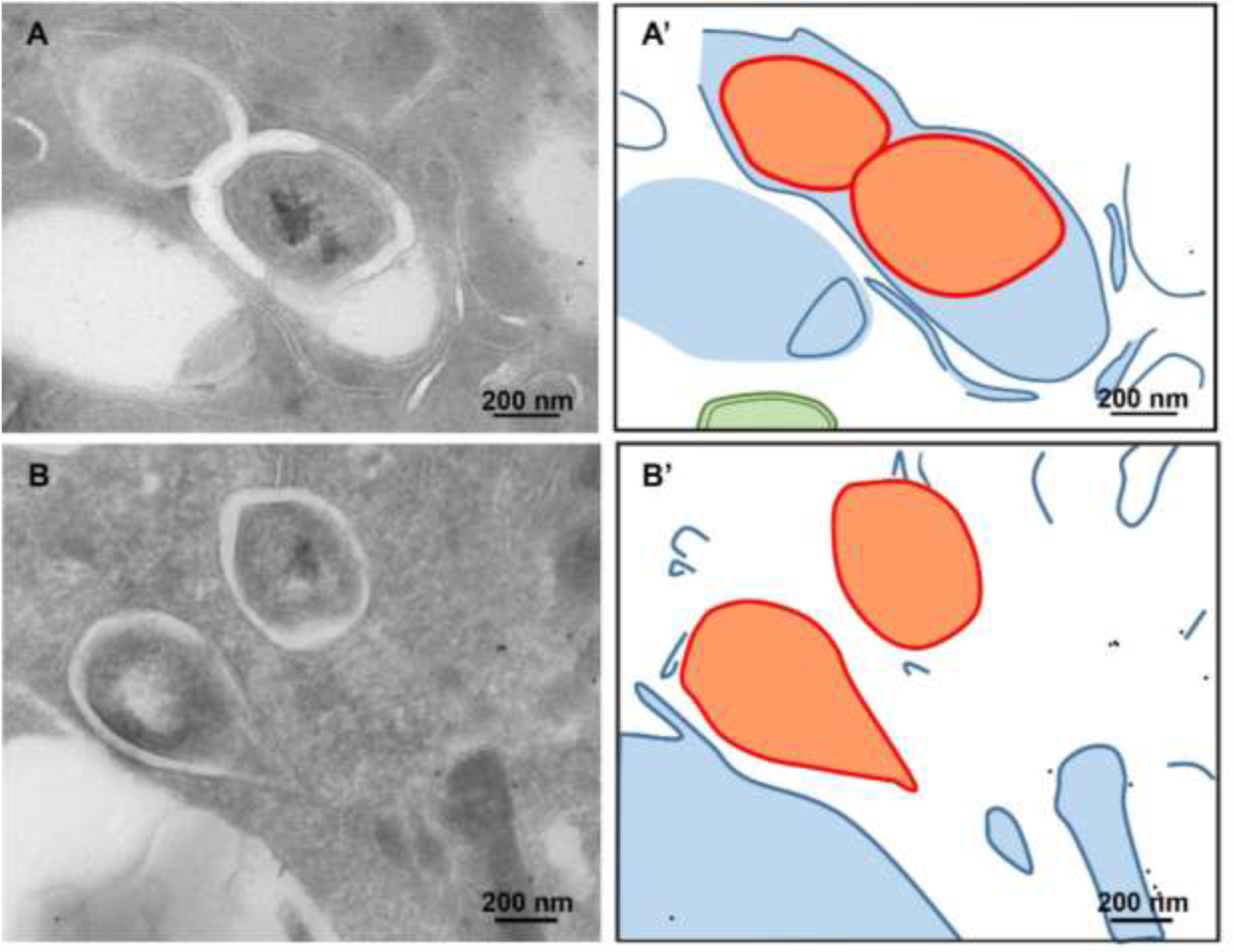
Early cytosolic localization of *Mtb* in SCID mouse lungs. A,B) Sections of SCID mice lung tissue infected with *Mtb* H37Rv for 21 days were labelled for LAMP1 using immuno-gold labelling, to indicate lysosomes. A’B’) schematic representation with in orange *Mtb*, blue lines host membranes, black dots gold particles indicating LAMP1 decorated lysosomes and green mitochondria. A) cross section of *Mtb* surrounded by cellular membranes thus phagosomal. B) *Mtb* is present in the cytosol, without cellular membranes surrounding the bacteria, scale bar indicates 200 nm. Information number of imaged bacteria and mice see Table S1.

To directly address a potential contribution of the adaptive immune system we next quantified *Mtb* cytosolic translocation in SCID mice. SCID mice lack functional T and B cells, key mediators of adaptive immunity in vertebrates and succumb rapidly to *Mtb* infections. SCID mice were infected with *Mtb* strain H37Rv via aerosol and sacrificed 21 days after infection. Lungs were then fixed and processed for EM analysis and sections labelled for LAMP1 using immuno-gold labelling to indicate lysosomes and phago-lysosomes (Table S1). We detected a 10-fold increase in *Mtb* cytosolic translocation in SCID compared to BALB/c mice, arguing that T and B cells are required for optimal clearance of cells with cytosolic bacilli.

### *Mtb* is preferentially located in the cytosol of infected pulmonary cells in IL-1R1 deficient mice

IL-1 is a potent innate inflammatory cytokine critically required for resistance against bacterial infections. Mice deficient in the IL-1 pathway, such as IL-1α and IL-1β are highly susceptible to *Mtb* infection with increased mortality and bacterial growth in lung and spleen and development of necrotic granulomatous lesions that more closely resemble human necrotic lesions (14, 15, 55–57). To investigate the contribution of this innate immune pathway in the prevention of mycobacterial cytosolic escape, we infected *Il1r1-/-* and B6 WT mice with *Mtb* and 4 weeks after infection fixed lung tissue and processed it for TEM analysis. As Mtb is normally difficult to find, first fluorescence microscopy was performed and tissues were sectioned at 200 nm and stained for detection of nuclei and mycobacteria (Fig. 6 A,B) to be able to select for the infected area (54). Whereas in B6 mice only small spots of *Mtb* are detected (like for BALB/c mice), infected *Il1r1-/-* tissue is heavily labelled and thus heavily infected. Ultrathin sections were labelled for LAMP1 and CD63 using immuno-gold labelling to indicate lysosomes and phago-lysosomes for TEM analysis. As before, difference between phago-lysosomal, phagosomal and cytosolic *Mtb* was determined based on the presence of immuno-gold labelling and a membrane surrounding the bacteria (Fig. 6C,D and Fig. S2D). *Mtb* was found in the phago-lysosome, the phagosome and in the cytosol. Strikingly, we detected a 14-fold increase in the number of *Mtb* translocated to the cytosol in *Il1r1-/-* deficient lungs compared to WT B6 lungs infected with *Mtb*. While only 2.5% of *Mtb* bacilli were located in the cytosol in the lungs of infected WT B6 mice, over 30% of the *Mtb* bacilli in the lung were able to escape to the cytosol in the absence of IL-1 signaling. Thus, IL-1 dependent signals are required to control cytosolic localization of *Mtb*.

**Legend Fig 6.**
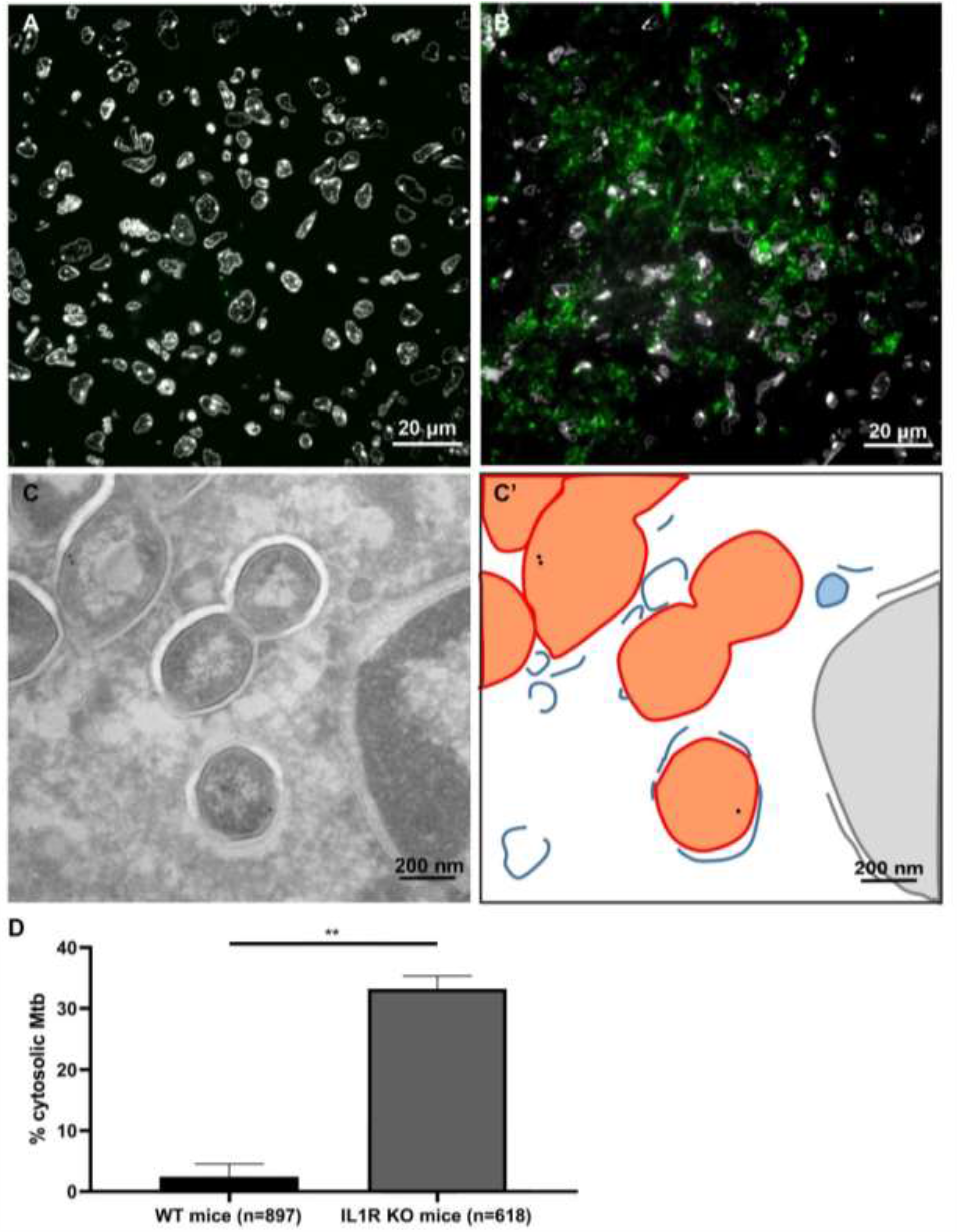
*Mtb* preferentially localize to the cytosol in *Il1r1-/-* mice. Fluorescence microscopy of 200 nm sections stained with DAPI for nuclei (white) and anti-cell wall protein to detect *Mtb* (green) in granuloma in lung tissue of A) WT B6 mice and B) *Il1r1-/-* mice infected with *Mtb* for 28 days. C) Immuno-gold labelling using LAMP1 indicate lysosomal membranes in *Il1r1-/-* mice imaged using TEM. C’) schematic representation of C, *Mtb* is depicted in orange, host membranes in blue and host nucleus in pink. D) Quantification of the localization of *Mtb* in B6 and *Il1r1-/-* lungs, here presenting cytosolic localization (see also Fig. S2D). The analysis included 897 (WT) or 618 (*Il1r1-/-*) bacteria and error bars indicate standard deviation based on in multiple granulomas of 2 WT B6 and 2 *Il1r1-/-* mice.

## Discussion

The cytosolic localization of mycobacteria has been debated ever since the first description in the late 1980s (18, 20, 58). Our immuno-gold TEM analysis of *Mtb*, BCG and mutant strains in 2007 (22) restarted the discussion and several studies have now confirmed the presence of extra-phagosomal *Mtb* bacilli (16, 21, 41). In addition, the release of DNA into the cytosol was described and is, like escape from the phagosomal compartment, dependent on the ESX-1 secretion system (29, 31, 32). In this study, we showed that *in vivo*, and with an intact immune response *Mtb* is mainly present in membrane bound compartments and not in the cytosol of infected cells. This also holds true for the pathogenic mycobacteria, *M. leprae* and *M. marinum*.

Here, we showed that raising the lysosomal pH did not affect cytosolic localization of *M. marinum*. In THP-1 cells, *M. marinum* can escape irrespective of the raised pH of the lysosome. As well-known already, maturation of the *Mtb* phagosome is altered (38, 44, 45) and our data suggest that the pH of phagosomes has no effect on translocation. Simeone and collegues (2015) (41) showed *in vivo* that when phagosomal acidification was blocked using bafilomycin, an induction of mycobacterial access to the cytosol is detected. Therefore, we hypothesized that raising the phagosomal and lysosomal pH by ConB would result in increased cytosolic localization of *M. marinum*. However, we found at a cellular level, no difference in the percentage of cytosolic *M. marinum*. Our cell-culture derived data merely demonstrate that mycobacteria can translocate irrespective of the pH, and thus manipulation of the pH by ConB did not change the subcellular localization in cells under *in vitro* conditions. Importantly, *in vivo*, as studied here and by Simeone et al., (2015) (41), subcellular localization of mycobacteria is a highly complex and dynamic process.

After inhalation in the lung, *Mtb* is initially taken up by alveolar macrophages and spreads among innate immune cells while delaying the initiating adaptive immunity (59–61). Importantly, both innate and adaptive immunity are critically important for optimal host resistance against *Mtb*. Here, we show that while the lysosomal pH itself has a minimal role, effective adaptive and innate host immunity play a critical role in regulating the number of cells with cytosolic mycobacteria. The delicate balance between bacterial replication and containment by the host immune response is lost when immunity is compromised by either lack of T and B cells in SCID mice, or even more dramatically, in the absence of innate IL-1R1 signaling. Of note, the profound increase in IL-1R1 deficient mice exceeds moderate but significant increase in SCID mice, even though the former likely has a broader set of immunological defects. This may argue that despite increased susceptibility to *Mtb*, certain specific immune pathways like IL-1 may be more directly and preferentially involved in regulating clearance of cells containing cytosolic bacteria. Activation of cytosolic immunosurveillance pathways, including those linked to IL-1, promote rapid clearance by the immune cells attracted to infected cells.

Cytosolic immunosurveillance pathways are triggered by pathogens directly or by pathogenic products entering the cytosol, and are often linked to anti-viral immunity and type I IFN induction as well as inflammasome activation. Inflammasomes are cytosolic signaling complexes that ultimately lead to cytolytic cell death and IL-1 family cytokine processing. It is now established that bacterial DNA can translocate to the cytosol, in an ESX-1 dependent manner, where it is detected by cytosolic DNA sensors such as cGAS and AIM2 (27, 29, 31, 32) cGAS in turn synthesizes a second messenger (cGAMP) which activates the STimulator of IFN Genes (STING) and type I IFN signaling and expression of IFN-α and β (reviewed in (62)). When cytosolic DNA is detected by AIM-2, the NRLP3 inflammasome is activated and leads to proteolytic cleavage of IL-1β (27, 29, 31, 32). Thus translocation of *Mtb* to the cytosol triggers a cascade of responses resulting at a cellular level in cell death (22, 63, 64). The connection between translocation, cellular damage and cell death has recently been studied in more detail; plasma membrane lysis caused by Mtb triggers inflammasome activation, IL-1β release and pyroptosis, a form of programmed necrosis (35). Beckwith et al (35) demonstrated that ESX-1-mediated phagosomal damage is a requirement for NLRP3 activation and although their live cell imaging demonstrated that inflammasome activation by *Mtb* is independent of lysosomal damage, others have demonstrated that active cathepsin release from ruptured phagolysosomes is a trigger of NLRP3 inflammasome activation (64). Taken together, after rupture of the phagolysomes by *Mtb*, cell death is induced and with an intact immune response, via the IL-1 signaling, these cells are cleared. This explains the fact that in skin biopsies of leprosy patients, armadillo, lung of BalbC mice, adult zebrafish the majority of the mycobacteria is detected in phagolysosomes.

We have previously shown that IL-1 and type I IFNs exhibit potent cross regulation important for host resistance against *Mtb* with excessive type I IFN induction in the absence of IL-1 signaling (14, 15, 65). The elevated type I IFNs expression in the absence of IL-1 contributed to the increased susceptibility of the *Il1r1-/-* mice, as mice doubly deficient in IL-1R1 and IFNAR1 displayed increased resistance (14). Our new findings here of increased cytosolic *Mtb* in *Il1r1-/-* mice provide a possible molecular explanation for the increased type I IFN production previously reported. In this context, our recent study demonstrated that infected cells themselves do not need to express IL-1R1 *in vivo* to mediate host resistance and that IL-1R1 expression coordinates immune responses in multiple cells types (13). Along these lines, it has been proposed that IL-1R1 on non-immune cells was required for the ability of infected alveolar macrophage to leave the airway to establish infection in the interstitial lung space (6). Thus, cytosolic containment of bacilli in infected cells, may not require direct cell-autonomous antimicrobial signaling pathways but may be the result of dynamic cellular interactions between infected cells and cells of both non-hematopoietic and bone marrow origin.

Overall, this study establishes that high level cytosolic escape of mycobacteria can indeed occur *in vivo*, but mainly when host resistance is compromised. When the complete adaptive immune system including B and T cells is abrogated, like in zebrafish embryos and SCID mice, a substantial percentage of mycobacteria is detected in the cytosol as compared to immunocompetent hosts. Strikingly, the highest proportion cytosolic *Mtb* was observed in mice lacking IL-1 signaling. This argues that the IL-1 pathway is crucial for the control of the number of cytosolic mycobacteria and likely IL-1 mediated resistance to *Mtb*.

## Methods

### Bacteria

*M. marinum* E11 strain was grown on Middlebrook 7H10 plates supplemented with OADC. A single colony was inoculated into 7H9 liquid medium (BD) supplemented with 10% ADC and 0.05% Tween 80 and incubated with shaking at 30°C and grown to an OD600 of 0.6-1. Before infection the bacteria were centrifuged at 750 rpm to remove clumps, leaving a bacterial suspension.

### Inhibiting acidification

Inhibition of acidification of phagosomes/lysosomes in THP-1 cells treated with Concanamycin B (ConB) (Alexis Biochemicals: 380098C100) was assessed by confocal microscopy. THP-1 macrophages were grown in Roswell Park Memorial Institute (RPMI)-1640 medium supplemented with 10% FCS and ConB was added to the cells in different concentrations 1h and 24h before fixation. The concentration 10nM of ConB strongly inhibited acidification and did not affect THP-1 cell viability after a 48h incubation period. THP-1 cells were washed 3 times with RPMI-1640 10% FCS medium and kept as a control or pretreated with 10nM ConB for 1 hour. Cells were infected with *M. marinum* (MOI 10:1) in the presence or absence of the acidification inhibitor. After an incubation time of 1 hour at 32°C, cells were washed 3 times with culture medium without antibiotics and with or without acidification inhibitors to remove extracellular bacteria. After washing, the cells were further incubated in culture medium with or without ConB for 24 or 48 hours at 32°C prior to fixation. Fixed samples were prepared for cryo-immunogold microscopy, sectioned for EM and immuno-gold labeled with CD63. To assess the effect on the amount of cytosolic *M. marinum*, from 3 independent experiments, 100-200 randomly chosen bacteria from one grid were counted and for each bacterium it was determined if it was cytosolic or resided in a phago-lysosome.

### DAMP assay for measuring lysosomal acidification

Luminal acidification in lysosomes was measured via the probe, DAMP [3-(2,4-dinitroanilino)-3′-amino-N-methyldipropylamine], incubated at 30 µM for 30 min to allow accumulation in acidic compartments. DAMP was quantified by immuno-gold staining using anti-DNP.

### Mouse infections

*IL1r1–/–* mice were purchased from Jackson Laboratories (JAX 3018) and backcrossed to C57BL/6 control mice from Taconic Farms (Hudson, NY) for 11 generations. Male and female mice, 8-12 weeks of age were infected via the aerosol route with *Mtb* H37Rv (100-200 CFU/mouse) as previously described (13) and sacrificed 27 days later. In short, mice were infected using a whole-body inhalation system (Glas-Col; Terre Haute, IN) exposing the mice to aerosolized *Mtb*. Lungs were perfusion-fixed in 4% paraformaldehyde and 0.4% glutaraldehyde overnight. After fixation, tissues were transferred to storage buffer containing 0.5 % paraformaldehyde. All animals were maintained in an Association for Assessment and Accreditation of Laboratory Animal Care (AALAC)-accredited BSL2 or BSL3 facilities at the National Institutes of Health (NIH) and experiments performed in compliance with an animal study proposal approved by the National Institute of Immunology Allergy and Infectious Diseases Animal Care and Use Committee.

### SCID Mice

Severe combined immunodeficiency (SCID) mice were purchased from Charles River and were aerosol infected with *M. tuberculosis* H37Rv for 21 days, when mice were sacrificed and lungs fixed by perfusion fixation as described below.

### BALB/c, B6 mice

Pathogen-free male BALB/c mice, 6-8 weeks old, were anaesthetized with sevoflurane vapours (Abbott Laboratories, Abbott Park, IL, USA) and 100 μl of PBS with 2.5 x 10^5^ viable H37Rv bacilli or either Beijing clinical isolates were inoculated intra-tracheally using a stainless steel cannula. Groups of 15 animals were then maintained in cages fitted with microisolators in a BSL-3 biosecurity level facility. Following infection, three mice were euthanized by exsanguination under anesthesia with pentobarbital at days 2, 7, 21, 45 and 120 of infection. Lung tissues were fixed by perfusion fixation as described below.

### Zebrafish

Zebrafish embryos and adult zebrafish (Danio rerio) were microinjected with *M. marinum* strain E11 as described by (53) at 30 °C. In short, zebrafish embryos were infected 28h post infection with 100 CFU *M. marinum* wild type through micro-injection in the caudal vein. Adult zebrafish were anaesthetized in 0.02% MS-222 (Sigma) and injected intra-peritoneally with 2×10^4^ *M. marinum* Tn::ESX-5. The embryos were incubated for 6 or 9 days and fixed as described below. For classification of the subcellular localization, 3 embryos of 9 days incubation were used. Three adult fish were infected with E11 or E11 ESX5 mutant (53) and sacrificed at day 11 when the spleen was dissected out and fixed as described below.

### Armadillo

Male Armadillos were intradermally inoculated in the abdomen with either 1 x 10^7^ live or irradiated *M. leprae* (0.1 ml). After, 3, 11 and 21 days post-inoculation, 6 mm punch biopsies were taken from the boarder of the depigmented area at the inoculation sites and fixed for 2 hours as described below. For classification of the subcellular localization, 1-2 punches were used from day 3 and day 21. As no lysosomal markers suited for immuno-TEM analysis were known, anti-Cell Wall Protein (CWP) was used to immuno-label the leprosy bacteria.

### Human samples

Leprosy skin biopsies were taken at the boarder of the depigmented lesions with written consent from 4 different lepromatous leprosy patients. Materials were directly incubated in EM grade fixatives and transported to the EM lab in fixatives.

### Fixation of tissues

All tissues were fixed in a combination of 4% paraformaldehyde and 0.4% glutaraldehyde in 0.2M PHEM (with 240mM PIPES, 100mM HEPES, 8mM MgCl2 and 40mM EGTA) buffer. Fixation for at least 2 hours in fixative containing glutaraldehyde is essential to kill mycobacteria. After fixation, tissues were transferred to storage buffer containing 0.5% paraformaldehyde in 0.2M PHEM buffer.

### Embedding and sectioning

After fixation, tissues were washed in phosphate buffered saline (PBS), to remove fixatives. The required structures were dissected out using a razor blade and cut into 1-3 mm^2^ sized blocks. Lung tissue was embedded in increasing percentages gelatin (2%, 5% and 12% in 0.1M phosphate buffer) and incubated at 37 °C. After the removal from liquid gelatin, blocks were incubated overnight in 2.3M sucrose at 4 °C. Then blocks were snap frozen and stored in liquid nitrogen till sectioned. After trimming at - 100 °C, semi thin sectioning was performed for analysis with fluorescence microscopy or ultra-thin sectioning at −120 °C was performed using a diamond Diatome cryo-immuno knife on a Leica Ultracut UC6. Sections are picked up with a loop filled with a 1:1 mixture of 2,3 M sucrose and 1% tylose (Metylcellulose, G1095) in milliQ water and placed on a copper, formvar coated grid or glass slides. Grids with sections were stored at 4 °C till immuno-labeled. Fluorescence Microscopy was used to search the region of infection as described in (54, 66). In short, semi-thin sections (200-300nm) of the whole sample were labelled with Hoechst 33342 (Thermo Fisher) to indicate the nuclei of the tissue and anti-Cell Wall protein labelling was used to indicate mycobacteria.

### Immuno-gold labelling

Grids with cryo-sections were incubated on 2% gelatin in 0.1M phosphate buffer plates at 37 °C for 30 min. Thereafter, grids were washed with PBS/0.02M glycine, blocked with 1% BSA and incubated for 45 minutes with primary antibody. Various antibodies were used on different tissues: for human skin and sputum: Cathapsin-B (Zymed, clone 1C11), Lysosome Associated Membrane Protein 1 and 2 (LAMP1 and LAMP2) (Pharmingen, H4A3 and H4B3), Cluster of Differentiation CD63 (clone 435 Sanquin). For Zebrafish anti-actin (Sigma AC-15), and tested but without specific labeling: anti-LAMP1 (Pharmingen, H4A3 and H4B3, Abcam ab67283), anti CD63 (clone 435 Sanquin). For *M. leprae*, anti Cell Wall Protein (C188, a kind gift form John Spencer and Patrick Brennan Colorado State University). DAMP (N-(3-((2,4-Dinitrophenyl) Amino)propyl)-N-(3-Aminopropyl) Methylamine, Dihydrochloride) detection was performed using anti dinitrophenol (DNP) (Polyclonal Anti-DNP, Oxford Biomedical Research). All antibodies were diluted in 1% BSA in PBS. After washing in PBS/0.02M glycine and blocking in 0.1% BSA in PBS/0.02M glycine, grids are incubated on bridging antibody when primary antibody was monoclonal or goat origin. After washing and blocking, grids are incubated on protein A gold diluted in 1% BSA in PBS. To remove unbound gold, grids were washed with PBS and fixed using 1% glutaraldehyde in PBS, then washed with MiliQ and contrasted with uranyl acetate and methylcellulose at pH 4.

### Immuno-fluorescence labelling

Semi-thin sections (200-300nm) placed on glass were washed with PBS/0.02M Glycine and incubated with primary antibody anti Cell Wall Protein for 45 minutes. Then, the sections were washed with PBS and incubated with secondary antibody goat-anti-rabbit Alexa 488 (Mol. Probes, A32731) for 20 minutes and Hoechst 33342 for 5 minutes (Thermo Fisher, H3570). After washing with PBS, samples were mounted with Vectashield. The sections were imaged using a Leica DM6 wide-field microscope and images were analyzed using FIJI.

### Statistical analysis and subcellular classification

Statistical analysis was performed using Graphpad-Prism 8.0 software. In the legends, the average is given with the standard deviation and n indicate the number of bacteria localized in a specific subcellular compartment. Significance was determined by using unpaired T test and defined in the graphs as P<0.05 = *, P<0.01 = ** and P<0.001 = ***. The number of biological samples, and bacteria counted in mice are listed in Table S1.

Classification of the subcellular localization of bacteria was performed blindfolded and by 2 individual counters to establish inter-counter reproducibility. Bacteria are classified as cytosolic when 1/3 or less of the bacteria or bacterial cluster is surrounded by a visible membrane and 2 or less gold particles detecting lysosomal markers are present. Bacteria are classified as phagosomal when 1/3 or more of the bacteria or bacterial cluster is surrounded by a visible membrane and 2 or less gold particles detecting lysosomal markers is present and phago-lysosomal when 1/3 or more of the bacteria or bacterial cluster is surrounded by a visible membrane and 3 or more gold particles detecting lysosomal markers are present.

### Ethics statement

SCID mouse infections were performed in agreement with European and French guidelines (Directive 86/609/CEE and Decree 87–848 of 19 October 1987). The experiments received the approval by the Institut Pasteur Safety Committee (Protocol 11.245) and the ethical approval by local ethical committees “Comité National de Réflexion Ethique sur l’Expérimentation Animale N° 59 (CNREEA)”.

Adult zebrafish of the local Free University of Amsterdam (VU) line were handled in compliance with the local animal welfare regulations and approved by the local animal welfare commission (IvD) of the VU/Amsterdam University Medical Centre. For zebrafish embryo experiments that are performed within the grace period (i.e. first 6 days) no special permission is allowed, since these experiments fall under animal experimentation law according to the EU Animal Protection Directive 2010/63/EU. BALB/c mouse infections were approved by the Institutional Ethics Committee of Animals Experimentation of the National Institute of Medical Sciences and Nutrition Salvador Zubirán in accordance with the guidelines of the Mexican national regulations on Animal Care and Experimentation NOM 062-ZOO-1999

Experiments using armadillos were performed in accordance with USPHS Policy on the Humane Care and Use of Laboratory Animals and the USDA Animal and Plant Health Inspection Service. The Institutional Animal Care and Use Committee reviewed and approved the protocol.

## Acknowledgements

We would like to thank Wikky Tigchelaar – Gutter, Pekka Kujala, Hans Janssen for EM analysis, Gidado Mustapha, Nigeria for leprosy skin biopsies. We are also grateful to Wafa Frigui and Alexandre Pawlik for help with infection experiments. We would like to thank Branch Moody for critically reading the manuscript and the helpful comments. RB acknowledges support by ANR-10-LABX-62-IBEID. SvdN PJP and NvdW acknowledge NIH grant no AI116604 and Netherlands Leprosy Relief. The NIH, NIAID funded the armadillo studies through the Interagency Agreement No. AAI15006 with the Health Resources and Services Administration, Healthcare Systems Bureau, National Hansen’s Disease Program.

## Author contributions

SvdN, ER, KDMB and NvdW wrote the manuscript, SvdN, MvZ, KdP, AG, SR, SM, DH, PP performed EM analysis AvdS, WB, RB, RHP, MP performed infection experiments, KDMB performed and designed experiments and NvdW conceived and designed experiments.

