## Supplemental Fig 1 for "IL-1R1 dependent signals improve clearance of cytosolic virulent mycobacteria *in vivo*"

**Supplemental Fig. S1**

**Fig. S1**

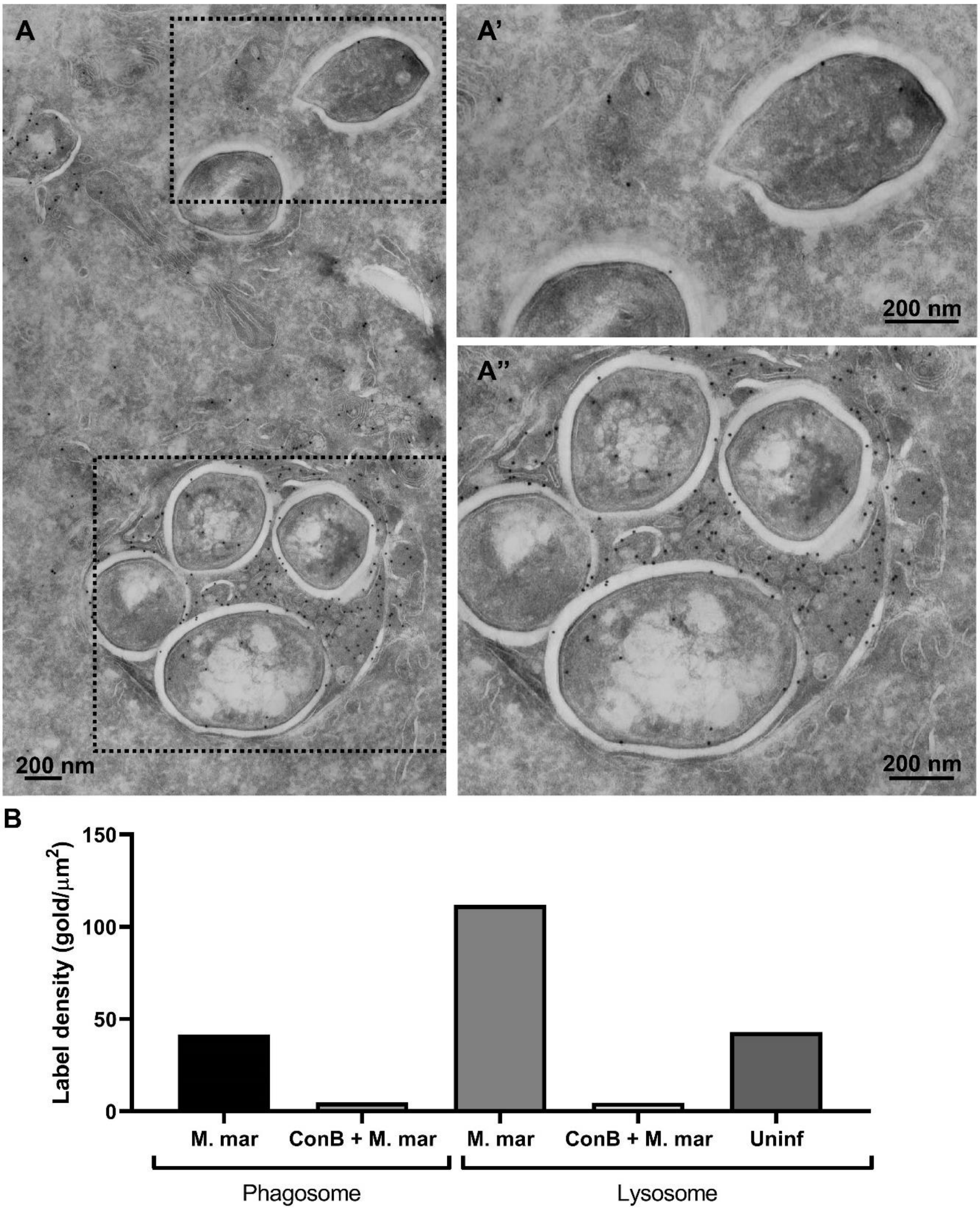

**Legend Fig. S1:**

The pH of the phago-lysosome does not affect cytosolic translocation of *M. marinum*. A) *M. marinum* infected THP-1 cells were incubated with DAMP to identify acidic organelles. Thereafter, the cells were fixed and analyzed using TEM. The acidity of the lysosome was determined using immuno-gold labelling against DAMP using DNP, the higher the label density the more acidic the phagosome or lysosome. A') cytosolic *M. marinum* without membranes enclosing the mycobacteria. A") Lysosomal *M. marinum*, enclosed by host membrane and DAMP labelled. B) Label density as measured in the number of gold particles per $\mu m^2$ on *M. marinum* containing phagosomes or lysosomes in *M. marinum* infected THP-1 cells, *M. marinum* infected Con B treated THP-1 cells and uninfected untreated THP-1 cells.
