## Supplemental Fig 2 for "IL-1R1 dependent signals improve clearance of cytosolic virulent mycobacteria *in vivo*"

**S Fig. 2**

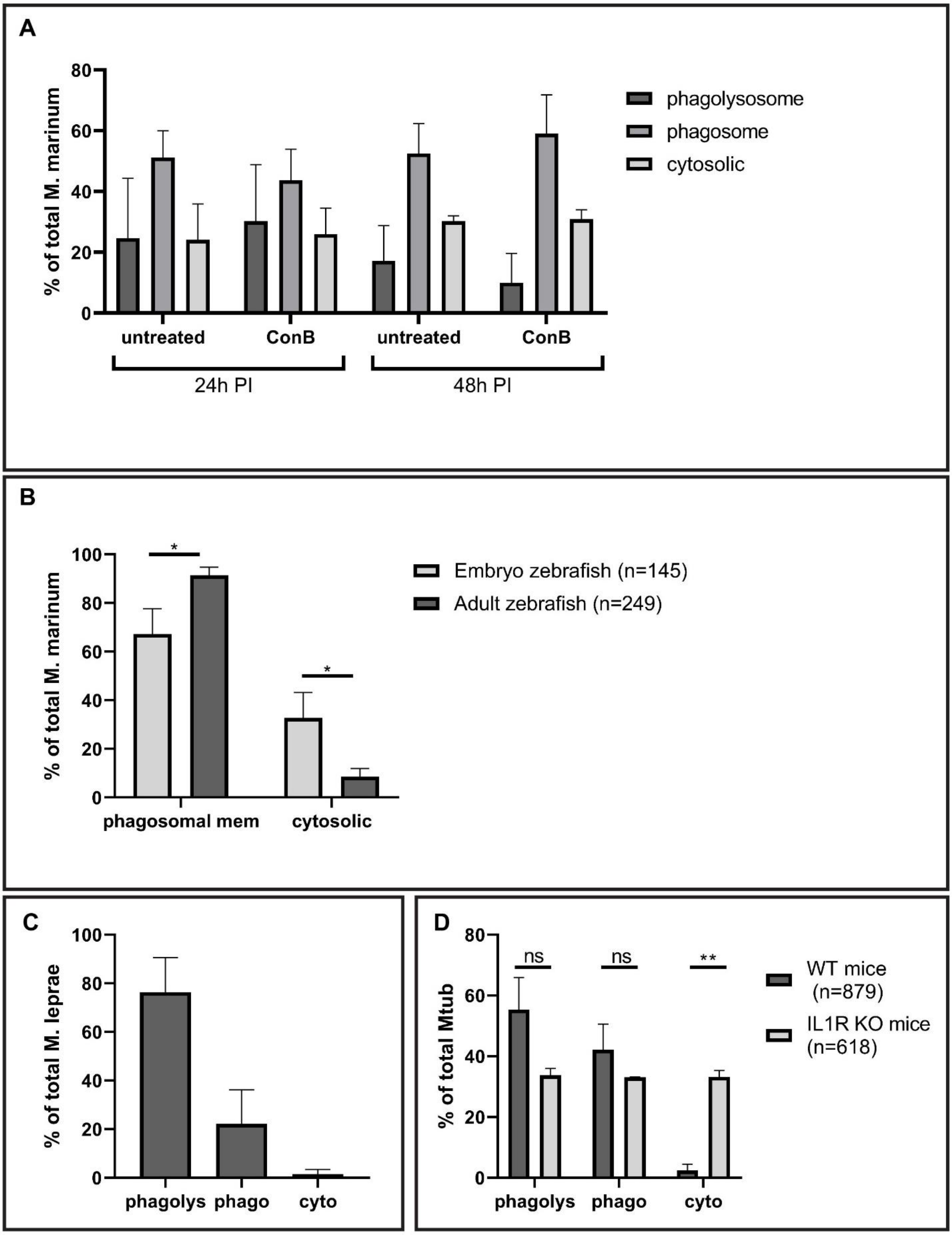
A
% of total M. marinum
0
20
40
60
80
phagolysosome
phagosome
cytosolic
untreated
ConB
untreated
ConB
24h PI
48h PI
B
% of total M. marinum
0
20
40
60
80
100
*
*
Embryo zebrafish (n=145)
Adult zebrafish (n=249)
phagosomal mem
cytosolic
C
% of total M. leprae
0
20
40
60
80
100
phagolys
phago
cyto
D
% of total Mtub
0
20
40
60
80
ns
ns
**
WT mice
(n=879)
IL1R KO mice
(n=618)
phagolys
phago
cyto

**Legend S Fig. 2**

**Quantification of the percentage of mycobacteria in the phagosome, phagolysosome or cytosol using immuno-gold labelling and TEM analysis.** The bacteria are classified as phagolysosomal when a host membrane is immunogold labelled with at least 2 gold conjugated to lysosomal markers such as CD63, LAMP1 or Cathepsin D are present, phagosomal with less than 2 gold marked membrane enclosed compartment are present and cytosolic when both membrane and gold is absent.

A) Subcellular localization *M. marinum* in THP1 cells treated with ConB or untreated. Immunogold labelling with CD63 average of 3 experiments with 100-200 bacteria classified (Related to Fig. 1B).

B) Subcellular localization *M. marinum* in zebrafish embryo day 9 and in spleen adult zebrafish day 11, error bars indicate standard deviation between 3 different zebrafish embryos and 3 adult fish. In three embryonic zebrafish, 17, 54 and 74 bacteria were detected and in three adult zebrafish 91, 8 and 150 bacteria were detected and catagorized. As no good lysosomal markers are present, a discrimination between phagolysosomal and phagosomal can not be made (Related to Fig. 2C).

C) Subcellular localization of *M. leprae* in skin biopsies of 4 different leprosy patients, using Cathepsin-D as a lysosomal marker, error bar indicates standard deviation from 4 different patients, where respectively 165, 248, 49 and 307 individual bacteria were detected and classified for their subcellular localization (Related to Fig. 4C).

D) Subcellular localization of Mtb in lung of WT B6 mice and Il1r1-/- mice 4 weeks infected using LAMP1 as a lysosomal marker (Related to Fig. 6D). Error bars represent standard deviation based on the analysis of 897 (WT) or 618 (Il1r1-/- ) bacteria in multiple granulomas of 2 WT B6 and 2 Il1r1-/- mice.
