## Supplement Table 1 for "IL-1R1 dependent signals improve clearance of cytosolic virulent mycobacteria *in vivo*"

**Table S1**

| Mouse | **Mtb strain** | **Day of infection** | **% bacteria phago-lysosome** | **% bacteria phagosome** | **% bacteria cytosol** | **n counted bacteria** | **n mice** |
|---|---|---|---|---|---|---|---|
| **BALB/c** | H37Rv | D2 | 38 | 62 | 0 | 47 | 1 |
| | | D7 | 71 | 23 | 6 | 52 | 2 |
| | | D21 | 0.4 | 98 | 2 | 247 | 2 |
| | | D45 | 0 | 100 | 0 | 5 | 1 |
| | | D120 | 0 | 97 | 3 | 38 | 2 |
| | 1998-1500 Ancient Beijing | D21 | 0 | 93 | 7 | 31 | 1 |
| | | D45 | 0 | 100 | 0 | 193 | 1 |
| | | D120 | 0 | 100 | 0 | 35 | 2 |
| | 2002-0230 Beijing | D120 | 0 | 100 | 0 | 166 | 2 |
| **SCID** | H37Rv | D21 | 29 | 54 | 17 | 167 | 1 |

**Legend Table S1:**

Overview of the subcellular localization of various *Mtb* strains infected at different days of infection in BALB/c or SCID mice. The number of bacteria used and the number of mice in which bacteria were detected are given in the last 2 columns. Bacteria are classified as phagolysosomal, phagosomal or cytosolic based on the number of gold attached to lysosomal markers and the presence of a membrane.
